# Structural Basis for Efficient Fo Motor Rotation Revealed by MCMD simulation and Structural Analysis

**DOI:** 10.1101/2025.06.28.662110

**Authors:** Shintaroh Kubo, Hiroyuki Noji

## Abstract

F_o_ domain of ATP synthase functions as a rotary molecular motor, coupling proton translocation with the rotation of the *c*-ring rotor. This process involves proton uptake at the entry half channel, rotor rotation, and proton release to the exit half channel. While the overall coupling mechanism is established, the design principle for efficient rotation remains unclear. Here, we employed hybrid molecular simulations—combining coarse-grained modeling and Monte Carlo methods—to investigate the roles of side chain flexibility at proton-binding residues and the angular mismatch between the proton uptake process and the proton release process. Our results indicate that both factors promote rotational activity, with side chain flexibility playing a more significant role. Comparable analysis of F_o_ structures from different species revealed that the key residue geometry is conserved, and that the asymmetric geometry of the two half channels aligns with the mechanism suggested by simulation. These findings highlight a conserved design principle that enhances rotational efficiency and offer a mechanistic basis for engineering synthetic rotary systems.

**Significance Statement:** F_o_ motor is the proton-conducting unit of ATP synthase and a rotary molecular motor driven by proton translocation across the membrane. Previous biochemical and structural studies have established the “half-channel model,” which explains how proton translocation is coupled with rotation. However, it remains unclear what structural features of F_o_ enable the smooth coordination of proton flow and rotation for kinetically efficient motion. In this study, we conducted a hybrid molecular simulation combining coarse-grained model calculations with Monte Carlo simulations. The present study found that the conformational flexibility of side chains in amino acid residues involved in proton translocation is one of the critical factors for kinetically efficient rotation. Furthermore, it is also found that the fundamental structural features are conserved across different species, suggesting that the principal mechanism and the design principles of F_o_ motor are well conserved across species.

## Introduction

F_o_F_1_ ATPase, one of the most ubiquitous enzymes responsible for the terminal reaction of oxidative phosphorylation, catalyzes the synthesis of ATP from ADP and inorganic phosphate, coupled with proton translocation driven by the proton motive force *(pmf*). This enzyme is composed of two rotary motors: F_o_ and F_1_ motors (Fig. 1a). These two motors are connected by a central rotor complex and a peripheral stalk. F_o_ motor is principally embedded in the lipid membrane, where it rotates the rotor ring relative to the stator complex of F_o_ upon proton translocation. The rotor ring is firmly bound to the rotary shaft of F_1_ motor, forming the central rotor complex in F_o_F_1_, while the stator complexes of F_o_ and F_1_ are connected by the peripheral stalk, so as to mutually transmit the rotary torque between F_o_ and F_1_. Under ATP synthesis conditions where *pmf* is sufficient, F_o_ motor generates larger torque and rotates the central rotor complex, inducing conformational transitions in F_1_ motor that results in ATP synthesis. When *pmf* is insufficient, F_1_ motor instead hydrolyzes ATP, driving the central rotor in the reverse direction and forcing F_o_ motor to pump protons.

**Figure 1.**
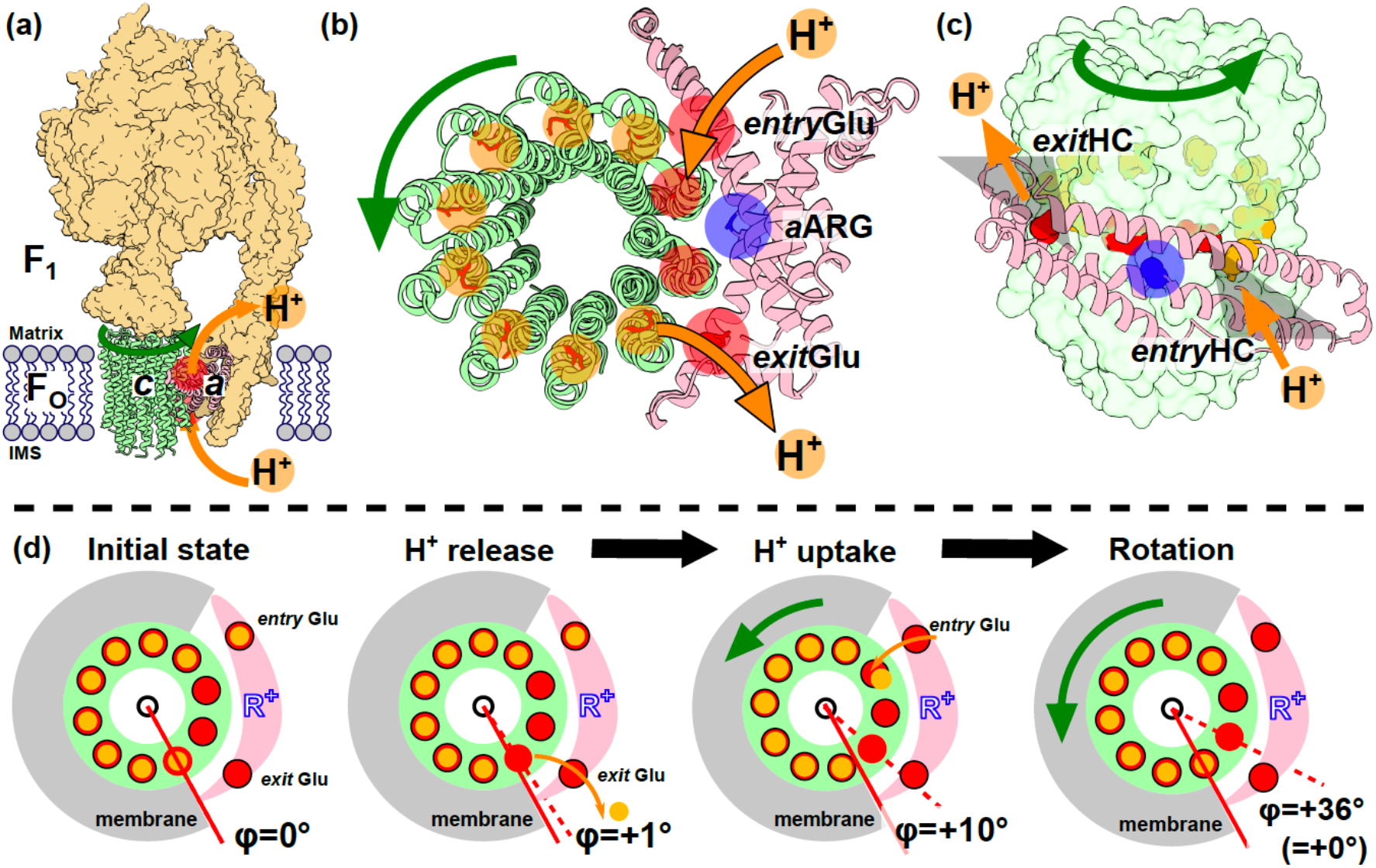
Rotational Mechanism of the F_o_ Motor. **(a)** Schematic representation of the overall model of F_o_F_1_ ATP synthase. F_o_ motor, including *c*-ring and *a*-subunit, is shown in cartoon representation. Green and orange arrows indicate the direction of the rotation and the proton translocation in ATP synthesis. **(b)** F_o_ motor viewed from the top. Half-transparent circles indicate the proton binding residues of the *c*-ring (*c*Glu) or the *a*-subunit (*entry*Glu, and *exit*Glu) in protonated state (orange) or deprotonated state (red). Blue circle indicates *a*ARG. **(c)** Viewed from the peripheral side. The two half-channels: *entry*HC and *exit*HC are depicted as half-transparent gray triangles. **(d)** Schematic of the proton transfer coupled with the rotation. Including *entry*Glu, and *exit*Glu, there are 12 protonatable glutamic acids. These are shown in orange when protonated, or in red when deprotoated. The initial angular position of the *c*-ring is defined as φ = 0°. Due to the 10-fold rotational symmetry, φ = 36° is equivalent to 0°.

F_o_ motor is principally composed of the *ab*_2_ stator complex and an oligomeric rotor complex of the *c*-subunit, termed the *c*-ring (Fig. 1b). The *c*-ring consists of multiple *c*-subunits arranged in a barrel-like ring. Each *c*-subunit consists of two membrane-spanning α-helices, with a highly conserved carboxyl residue (generally glutamate, *c*Glu) located midway along the N-terminal helix; this residue serves as the proton-binding site. The *a*-subunit contains two membrane half-spanning channels at the interface with *c*-ring, so-called *‘half channels’*, which comprise part of the proton-conducting pathway of F_o_. When viewed from the side of the membrane, each half-channel extends from a central midpoint to one of the aqueous compartments (Fig. 1c), and the two half-channels traverse the membrane in opposite directions. Hereafter, the half-channel that opens to the periplasmic space in bacteria or the inner membrane space in mitochondria is referred to as “*entry*HC”, since F_o_ uptakes protons there during ATP synthesis. The other half-channel, “*exit*HC”, is where protons are released into the cytoplasm (in bacteria) or the matrix (in mitochondria). Viewed from the top (Fig. 1b), these two half-channels lie at distal positions from each other and are separated by a highly conserved arginine residue (*a*Arg).

Structural data and biochemical experiments support the *half-channel model* for the rotation of *c*-ring rotation coupled with proton translocation. In this mechanism, protons are translocated across the membrane via two proton-transfer (PT) events — one between the conserved glutamate in *entry*HC (*entry*Glu) and *c*Glu, and another between the conserved glutamate in *exit*HC (*exit*Glu) and cGlu. During ATP synthesis, starting with proton uptake, a proton in the periplasmic space of bacterial or inner membrane space of mitrochondria enters *entry*HC, protonating *entry*Glu. The proton is then transferred to *c*Glu of the adjacent *c*-subunit (entry proton transfer, *entry*PT) (Fig. 1b and 1c), neutralizing the negative charge of *c*Glu allowing the *c*-subunit to be exposed to hydrophobic environment of lipid bilayer. This induces the rotation of the *c*-ring while the other proton transfer is also requisite for the rotation as mentioned below. After nearly one full turn, the proton-carrying *c*-subunit aligns with *exi*tHC, where the proton is transferred to *exit*Glu (exit proton transfer, *exit*PT) and released to the opposite side of the membrane.

In this model, the *c*-ring rotation is biased in the counterclockwise direction by two “ratchet” mechanisms. One is the deprotonated *c*Glu at *exit*HC which prevents backward rotation owing to the high energetic penalty of exposing its negative charge to the lipid environment. The second ratchet is the gate-keeper residue *a*ARG: this positively charged arginine recognizes deprotonated *c*Glu, permitting rotation, but disallows rotation when *c*Glu is protonated. Thus, *a*ARG prevents protons from directly moving between *entry*HC and *exit*HC without being coupled with the *c*-ring rotation (1–6). From the standpoint of the *c*-ring, an individual rotation step begins after these two proton-transfer events (Fig. 1d).

The number of *c*-subunits ranges from 8 to 15, depending on the species. In the yeast mitochondria and some bacteria (7–9), the *c*-ring is composed of ten *c*-subunits and is expected to complete one full rotation in ten steps, each step coupled to the translocation of a single proton (for a total of ten protons per rotation). A previous study employing Fluorescence Resonance Energy Transfer (FRET) observed *c*_10_-ring rotation and identified ten frequently visited “hotspots” (10), consistent with the above model. In contrast, we observed three clear clear steps per turn under ATP-synthesis conditions (11), suggesting that the apparent step size is determined by whichever motor — F_o_ or F_1_ — is rate-liming. While these stepping behaviors in F_o_F_1_ ATP synthase have been characterized, the detailed mechanism by which rotation couples with proton translocation remains elusive.

Coarse-grained molecular dynamics (MD) simulations have successfully revealed detailed conformational transitions and energy profiles for diverse molecular motors and complexes, including F_o_F_1_ ATPase (12–17), MotAB (18), and Cytoplasmic Dynein (19–22). This approach allows for the simulation of large-scale conformational dynamics of molecular motors and molecular machine complexes, which is practically infeasible with all-atom MD simulations.

In a previous study, we have developed a hybrid Monte Carlo/Molecular Dynamics (MC/MD) simulation method to investigate the coupling between rotational motion and proton translocation in F_o_ motor (12). Because the timescale of proton transfer (PT) is much shorter than that of conformational dynamics, residues involved in PT (exclusively glutamates in this system) were treated as being either protonated or deprotonated, neglecting intermediate states. In this model, the probability of PT was calculated from the system’s energy difference before and after PT. Then, the probability was used in the MC procedure. When this method was applied to the F_o_ motor with *c*_10_- ring rotor from the yeast mitochondria, ten low-energy regions was reproduced, consistent with the ten-fold structural symmetry of the rotor (12). The simulations also showed that in each round of rotation step, the two PT events do not occur in a random order, but in a biased sequence: first at *exit*HC, then at *entry*HC after a slight rotational shift. These findings are corroborated by other studies (23–25).

A recent cryo-EM study has revealed that, in addition to the stable conformational state universally observed in F_o_F_1_, there is a sub-stable state in which the *c*-ring is rotated by about 10° from the usual stable position (26). Consistent with this, all-atom MD simulations identified a corresponding sub-stable state near 10° (23–25). These simulations suggest that when the *c*-subunit facing *exit*HC releases a proton to *exit*Glu (*exit*PT state, ‘H^+^ release’ in Fig. 1d), the resultant conformation corresponds to the 0° stable state found in most of reported structures. On the other hand, once the *c*- subunit adjacent to *entry*HC uptakes a proton from *entry*Glu (*entry*PT state, ‘H^+^ uptake’ in Fig. 1d), the *c*-ring shifts by about 10°. These findings indicate that *entry*PT and exitPT occur at distinct rotational angles of the *c*-ring, implying that the angular mismatch may be significance for F_o_’s efficient rotation.

In this study, to gain deeper insights into F_o_’s design principle, we first ask whether the angular mismatch between *entry*PT and exitPT is important. Such structural mismatch might be compensated by conformational flexibility of the residues involved in the proton transfer events. Hence, we also ask whether that flexibility plays a discernible role in *c*-ring rotation. In particular, we focus on the flexibility of the proton-carrying residue *c*Glu, as the carboxy group of the side chain has been observed to shift conformation even though the overall *c*-ring structure remains relatively rigid. To address these questions, we have applied our previously developed hybrid MC/MD simulation framework (12) to simulate *c*_10_-ring of yeast mitochondrial F_o_. We then performed a comprehensive survey of F_o_ in the light of our simulation results. This combined approach offers new insights into the general operational and design principles of F_o_.

## Results

### Hybrid MC/MD simulation

The hybrid MC/MD simulation system comprised the *c*_10_-ring and *a*-subunit of F_o_ from *S. cerevisiae* mitochondria, as in our previous study (12)(Fig. 2a). The reference structure for the simulation was PDB ID: 6CP6 (7). In this system, *c*Glu, *entry*Glu and *exit*Glu correspond to E59 of the *c-*subunit (*c*E59) and E223/E162 of the *a*-subunit (*a*E223 and *a*E162), respectively. For the MD part of the simulations, every amino acid residue was represented by a single sphere of which diameter reflected its effective size-including the size chain-, following a coarse-grained model (27,28). Acidic and basic residues were assigned charges of -1 or +1, respectively. The three key residues for proton transfer (PT) — *c*Glu, *entry*Glu, and *exit*Glu —was modeled as either protonated (uncharged) or deprotonated (−1 charged). Their protonation states were updated using the Monte Carlo (MC) scheme. The main MC protocol is outlined below; full details are given in our previous report (12). In each MC update cycle, we first evaluated the protonation states of *entry*Glu and *exit*Glu, taking into account their pKa values and the effective local pH in *entry/exit*HC (calculated from the applied *pmf*, 150 mV). We then determined the protonation state of *c*Glu, computing the probability of PT between *entry/exit*Glu and *c*Glu based on the electrostatic potential of the system and the effective distance between the C*a* atoms of the paired residues. The PT probability was set as zero when both *entry/exit*Glu and *c*Glu are protonated or deprotonated. Because our coarse-grained model does not explicitly include side chains, the exact positions of the carboxyl groups cannot be used directly to define the effective PT distance. To compensate, we introduced a *virtual vector* originating from the C*a* of *c*Glu (dark green arrows in Fig. 2b) to represent the side chain orientation. The default orientation of this vector was set to 60° relative to the tangent line of the *c*-ring arc, as observed in the reference structure; this orientation was set as 0°. The effective PT distance was thus calculated from both the C*a-*C*a* distance and the orientation of the virtual vector. After every 10^5^ MD steps in the MD simulation, MC update was executed, after which the next round of MD segment began. This MC/MD cycle was repeated 30 000 times (30 000 frames) to generate each trajectory.

**Figure 2.**
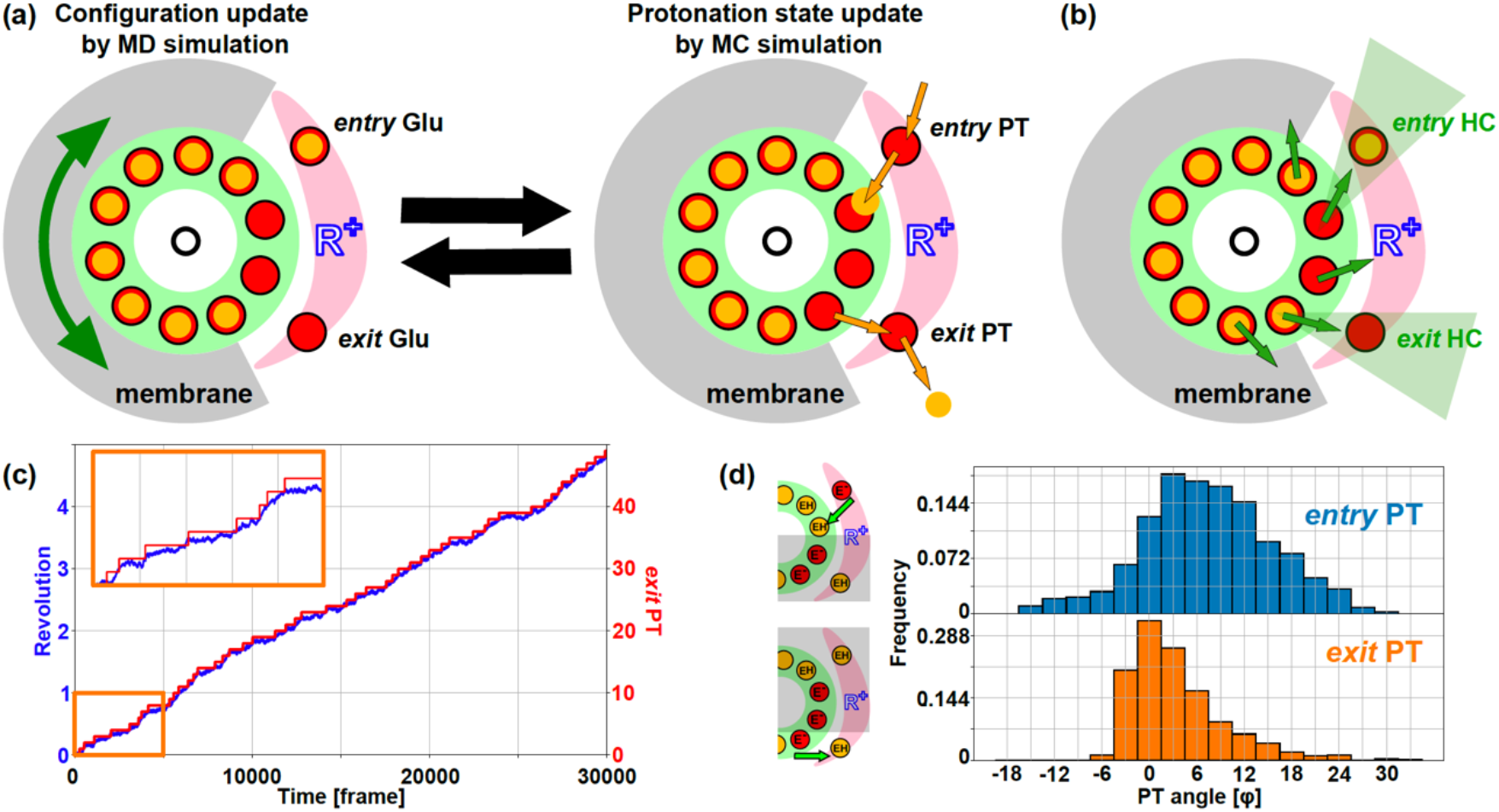
Relationship between proton translocation and rotation. **(a)** Flowchart of the MC/MD simulation. **(b)** Illustration of the “virtual vector” represents effective orientation of carboxy group of *c*Glu’s. **(c)** Trajectories. The blue shows the rotational angle of *c*-ring, and the red shows the cumulative number of protons exiting to the *exit*HC. **(d)** The histograms of the angular position of *c*-ring for proton transfer (PT): stator-to-rotor PT at *entry*HC (*entry*PT) (blue), and rotor-to-stator PT at *exit*HC (*exit*PT) (orange). The angle φ is defined as the deviation of the rotational angle from the initial position in the synthesis direction (counterclockwise).

### Rotation after proton exit

The MC/MD simulation showed that *c*-ring rotates in the synthesis direction with net proton translocation under *pmf* of 150 mV (Fig. 2c). In each round of 36° step rotation, the two PT events: *entry*PT and *exit*PT were observed, translocating a net of a single proton across membrane. This is evidently shown in Fig. 2c where *exit*PT events were followed by the rotation. As reported previously, the two PT events were principally stochastic but occurred in the biased sequence: *exit*PT occurred first, followed by *entry*PT.

Fig. 2d shows the histograms of the *c*-ring’s angular positions for *entry*PT and *exit*PT occurs. Although these PT events span a broad range of angles and overlap, it is evident that the mean angles for the two events differ. *exit*PT occurs most frequently at φ = 0°, whereas *entry*PT often occurs after a slight rotation of approximately 3.6°. This finding can be explained by the asymmetric positioning of *exit*Glu and *entry*Glu relative to the membrane: *exit*Glu is located closer to the membrane. Its proximity imposes a higher energy barrier on the negatively charged *c*Glu at the *exit*HC following *exit*PT, inducing the slight rotation. In contrast, *c*Glu at the *entry*HC does not encounter such a high energy barrier, which allows *entry*PT to occur in a broader range of angles.

### Effect of virtual vector on rotation activity

From the observation of the slight rotational motion after *exit*PT, we considered the possibility that the rotational shift plays a role as a remote interplay between the two PT events for efficient rotation, *i. e*. positive allosteric effect. Therefore, considering that the difference between the angular positions for *entry*PT *and exit*PT is key for smooth rotation, we hypothesized angular mismatch for *entry*PT *and exit*PT would enhance rotation velocity. To modulate the angular mismatch with the minimum change in the MC/MD simulation model, we varied the orientation of virtual vector (ψ) of *c*Glu at *entry*HC while the virtual vector of *c*Glu at *exit*HC remained unchanged. In addition, we considered that the rotational fluctuation of the virtual vector (σ) is another key factor for the rotation velocity. In the default simulation set-up above, the virtual vector had rotational distribution from the C*a* of *c*Glu as a Gaussian function with ψ = 0° and σ = 36°. In followings, we aimed to investigate the effect of these factors on *c*-ring rotational velocity and the angular distribution of proton translocation by systematically varying ψ for *c*Glu at *entry*HC and σ for both of *c*Glu’s at *entry*/*exit*HC (Fig. 3a).

**Figure 3.**
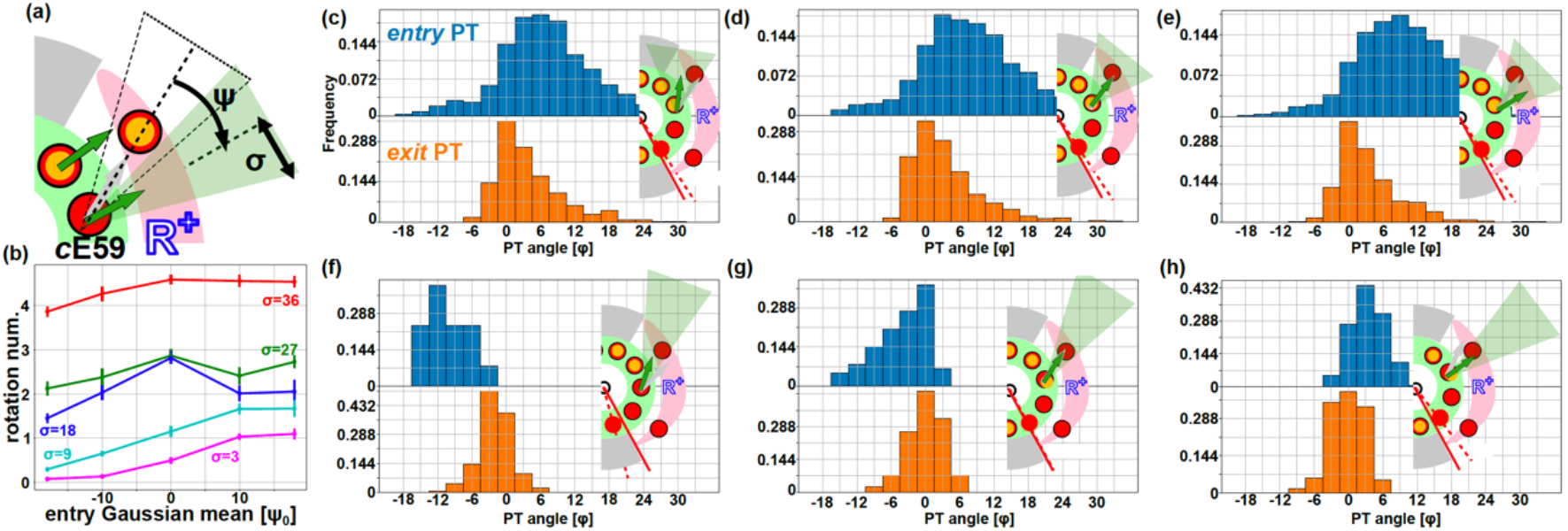
Changes in rotational activity with entry θ_0_. **(a)** Schematic representation of the mean value (ψ) and standard deviation (σ) of the virtual vector. When ψ is positive, the orientation of virtual vector is rotated clockwise. While the ψ was changed for *c*Glu at *entry*HC, σ was changed for both *c*Glu at *entry*HC and *c*Glu at *exit*HC. **(b)** The mean cumulative rotation number versus ψ with different σ conditions: σ=36 (red), 27 (green), 18 (blue), 9 (cyan), 3 (magenta). **(c-e)** Histograms of *entry*PT and *exit*PT frequencies against the *c*-ring rotational orientation (φ) when σ=36° (**c**; ψ=-18°, **d**; ψ_0_=0°, **e**; ψ_0_=+18°**). (f-h)** Histograms when σ=3° (**f**; ψ=-18°, **g**; ψ_0_=0°, **h**; ψ_0_=+18°).

Fig. 3b summarizes the results. There are two obvious trends observed. Firstly, larger σ results in faster rotation. With σ = 36°, the cumulative number of rotations in the simulation time (30k frames) exceeded 4 rotations in most conditions, whereas the smaller σ conditions (3° or 9°) yielded only 1-2 revolutions at most. This trend is reasonable, considering that larger σ enables proton transfer in a wide range of angular position of *c*-ring that allows smooth rotation. The second trend is that a larger ψ is also effective for rotation. This trend is more evident when σ is small (9° and 3°), where the virtual vector fluctuation is restricted. Although the angular mismatch enhanced rotation velocity, the effect is asymmetric; positive ψ enhanced rotation, while negative ψ suppressed rotation. In addition, the effect of positive ψ is not significant when the side chain fluctuation is large, *i. e*. at larger σ values.

To explore the mechanism of these phenomena, PT frequency was plotted against the rotation angle, φ (Fig. 3c-h, Supplemental Fig. 1a). When σ is large, the PT frequency was broadened as expected (see Fig. 3c, d, e for σ of 36°). Interestingly, the distribution did not change with ψ. This is consistent with the almost constant rotation activity, around 4 revolutions, irrespective of ψ values. The angle difference between *entry*PT and *exit*PT were also conserved; the main peaks for *exit*PT are found at around 0° while that for *entry*PT are found between 3-9°. These observations suggest that the side chain fluctuation masked the effect of ψ, and that the principal sequence of the reactions is conserved; *exit*PT occurs first, followed by slight rotation and *entry*PT, although the angle difference between *entry*PT and *exit*PT slightly increased from 3° to 9°.

On the other hand, when σ is small, the PT frequency is sharpened and the effect of ψ becomes remarkable (Fig. 3f, g, h). While the main peak position for *exit*PT remains around 0°, that for *entry*PT is shifted minus when ψ=-18° and shifts plus when ψ=+18°. Because ψ for *c*Glu at *exit*HC was kept at the original angle (0°), the *exit*PT occurs in principle before *entry*PT. Therefore, the peak shift of *entry*PT toward the minus direction means that the *c*-ring reversibly rotated to initiate *entry*PT. This is reasonable because when the virtual vector is oriented in minus (ψ is negative), the position of *c*Glu for *entry*PT has to be reversibly rotated to adjust the virtual vector orientation toward *entry*Glu. However, the *c*-ring is principally pushed in the forward direction upon *exit*PT. This is because *exit*PT results in the potential slope for *c*-ring due to the energetic penalty against the deprotonated *c*Glu in proximity to the membrane environment (12). Therefore, the *c*-ring has to be reversed against the potential slope for *entry*PT, that is the suppression mechanism for the rotation when ψ is negative. On the other hand, when ψ=+18°, the peak for *entry*PT is shifted forward and the rotation activity is enhanced. This is also attributed to the slight rotation of *c*-ring after *exit*PT; upon the slight rotation, *c*Glu for *entry*PT can orientate its virtual vector to *entry*Glu.

Thus, the peak angle mismatch for *entry*PT and *exit*PT correlates with rotation activity at low σ conditions. For a comprehensive analysis, we plot the rotation activity against the peak angle difference (*entry*PT-*exit*PT) (Supplemental Fig. 1b, c) of all conditions tested. The plot shows a clear correlation between the rotation activity and the peak angle difference, suggesting the peak angle mismatch caused by the rotational drift after *exit*PT is a good barometer for rotation activity. As discussed above, the rotational drift is induced by the energetic penalty against the deprotonated *c*Glu after *exit*PT in proximity to the membrane environment. Taking this into account, the key structural basis for the rotation activity can be attributed to the proximity of *exit*Glu to the membrane, the fluctuation of the side chain of *c*Glu, and the angular offset between the two PT events at the exit and entry sites.

### Comparative analysis of F_o_ structures

In the above hybrid MC/MD simulations, we analyzed F_o_ from yeast mitochondria. To evaluate whether the identified mechanisms for efficient rotation are broadly applicable and to gain new insights, we conducted a comparative analysis of F_o_ structures from other species that have a different number of *c*-subunit in the *c*-ring; *c*_8_-ring (29–31), *c*_9_-ring (32), *c*_10_-ring (7,9,26,33,34), *c*_12_-ring (35), and *c*_14_-ring (36) (Table 1).

**Table 1.** cN-ring PDB list.

| cN | Origin | PDB | entry | aARG | exit | cGLU |
| --- | --- | --- | --- | --- | --- | --- |
| 8 | *Bovine* | 6zpo | E203 | R159 | E145 | E58 |
| 8 | *Human* | 8h9s | E203 | R159 | E145 | E58 |
| 8 | *Sus scrofa* | 6j5i | E203 | R159 | E145 | E58 |
| 9 | *Mycobacterium Smegmatis* | 7jg5 | D222 | R188 | E177 | E65 |
| 10 | *Bacillus PS3* | 6n2y | E178 | R169 | E159 | E56 |
| 10 | *E. coli* | 6oqr | E219 | R210 | E196 | D61 |
| 10 | *S. cerevisiae* | 6cp6 | E223 | R176 | E162 | E59 |
| 10 | *Pichia augusta*<br>(*Ogataea angusta*) | 5lqz | E226 | R179 | E165 | E59 |
| 10 | *Polytomella* | 6rd9 | E288 | R239 | E225 | E111 |
| 12 | *Paracoccus dentrificans* | 5dn6 | E1256 | R1217 | E1203 | E60 |
| 14 | *Spinacia oleracea* | 6fkf | E198 | R189 | E178 | E61 |

Fig. 4a-c shows the representative structures: F_o_ with *c*_10_-ring from *S. cerevisiae* mitochondria, F_o_ with *c*_8_-ring from *Bovine*, and F_o_ with *c*_14_-ring from *S. oleracea*. We first examined three structural features: the central angle of *c*-ring sector between *entry*Glu and *exit*Glu (Fig. 4d), the triangle formed by three key residues (*entry*Glu, *exit*Glu, and *a*ARG) (Fig. 4e), and the number of *c*-subunits covered by *a*-subunit (Fig. 4f). The reference F_o_ with *c*_10_-ring from *S. cerevisiae* mitochondria exhibits the central angle of 68.9°, and the coverage number of 3.76. Other F_o_ structures show similar central angles (around 63.4), regardless of the number of *c*-subunit or species. The coverage number was also similar across species, around 3.4, except for *c*_9_-ring (2.4) and *c*_14_-ring (3.7). The triangular configuration of three key residues was also broadly conserved though species with a larger number of *c*-subunit (e. g. c_12_, and c_14_) show the reverse orientation of the triangles, likely a broader arc of *a*-subunit associated with the larger radius of *c*-ring.

**Figure 4.**
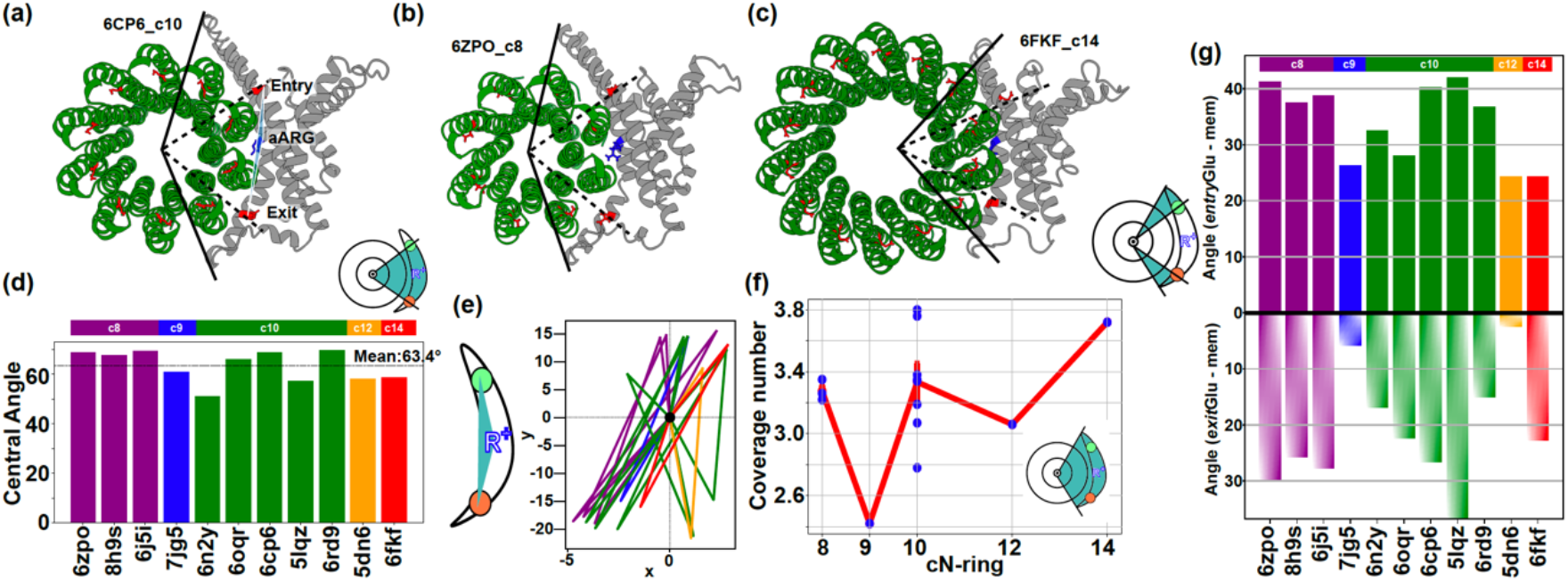
Structural comparison of cN-ring. **(a-c)** F_o_ motor from PDB ID: 6CP6, *c*_10_-ring **(a)**, 6ZPO, *c*_8_-ring **(b)**, and 6FKF, *c*_14_-ring **(c)**. The *c*-ring is shown in green, and the *a*-subunit in gray. The light blue triangle is composed of three points: *a*ARG (blue), *entry*HC (red), and *exit*HC (red). The black line is the a-subunit angle that formed between the center of the N proton-transporting acidic residues of the *c*-ring and the maximum and minimum y-axis values of *a*-subunit. Also, the dashed black line is the angle between the entry and exit. **(d)** The angle between the entry and exit plotted by dashed black line list. **(e)** Composed light blue triangle as shown in **(a)**. The coloring is same with **(d). (f)** Number of c-subunits covered by *a*-subunit. Blue circles represent the corresponding positions for each PDB, and the red line shows the average values and standard errors for each N. **(g)** The angle between the top of *a*-subunit and entry (upper), and the bottom of *a*-subunit and exit (under). The coloring is same with **(d)**.

Interestingly, in all species analyzed, the distances from *a*Arg to *entry*Glu and *exit*Glu were consistently asymmetric, with *entry*Glu positioned closer to *a*Arg (Fig. 4e). What functional implication does this asymmetry have?

This feature can be rationalized in the context of the MC/MD-derived mechanism: after proton release from *c*Glu to *exit*HC, the *c*-subunit becomes energetically less stable near the hydrophobic membrane and is pushed toward *a*Arg. Upon forming an electrostatic interaction with *a*Arg, the *c*-subunit must then overcome this interaction thermally to rotate further and undergo proton uptake at the entry site (*entry*PT). The proximity of *entry*Glu to is positioned closer to *a*Arg than *exit*Glu may thus facilitate the transition, supporting efficient rotation.

This suggests that the proximity of *exit*Glu to the membrane environment may also be a key feature for rotation activity. We therefore examined the relative proximity of *exit*Glu and *entry*Glu to the membrane. As shown in Fig. 4g, *exit*Glu was consistently located closer to the membrane than entryGlu in all species analyzed. This asymmetry was particularly pronounced in *c*_9_- and *c*_12_-ring F_o_ motors—those the number of *c*- subunits divisible by three. While the underlying mechanism for this feature remains unclear, it is a notable and potentially important observation.

Taken together, these findings demonstrate that the structural features identified in yeast F_o_ are largely conserved across species and *c*-ring sizes. The asymmetric positioning of *entry*Glu and *exit*Glu relative to *a*Arg aligns well with the biased rotation mechanism observed in simulations. Moreover, the consistent proximity of *exit*Glu to the membrane further supports the generality of the simulation-derived mechanism. These results suggest that the mechanistic features elucidated through the yeast F_o_ model likely represent a broadly conserved design principle. Future studies should aim to validate this conservation through comparable hybrid simulations. Additionally, the structural variations observed in some F_o_ motors warrant further investigation into how they influence the coupling mechanism between proton translocation and rotary motion.

## Discussion and Conclusion

In this study, we investigated the angular positions at which two proton transfer occur—either after proton uptake at the entry half-channel (*entry*HC) or before proton release at the exit half-channel (*exit*HC)—and how the distribution of these events affects the rotational velocity of the *c*-ring. Using MC/MD simulations, we systematically varied the width and mean orientation of a virtual vector representing the side chain orientation of the glutamic acid on the *c*-subunit (*c*Glu) to evaluate the influence of these parameters on rotation. Our results showed that rotational velocity increased when the virtual vector fluctuates in a wide range or when the virtual vector for *entry*PT was oriented backward. It was also shown that the vector fluctuation has more significant effect. Comprehensive analysis of proton transfer angle showed a clear correlation between the angle mismatch between *entry*PT and *exit*PT and the rotation activity. These findings underscore the importance of allowing proton transfer to occur over a relatively broad angular range.

While *c*-rings exhibit different symmetries across species, comparative structural analysis of 11 F_o_ complexes revealed that the spatial relationship among three key residues on *a*-subunit—*entry*Glu, *a*ARG, and *exit*Glu—remains largely conserved. This suggests that the system does not finely tune the positions of these sites according to the number of *c*-subunits, but instead adjusts using the flexibility of the side-chain orientations. This structural consistency supports the simulation-based conclusion that a broad angular range for proton translocation is essential for efficient rotation.

Interestingly, all analyzed structures showed that *a*-subunit consistently interacts with about 3 to 4 *c*-subunits, regardless of *c*-ring’s symmetry. For example, in *c*_8_, *a*- subunit appears to cover a broader arc, whereas in *c*_14_, it spans a narrower region—but the actual number of interacting *c*-subunits remains similar. This suggests that key electrostatic interactions between *a*-subunit and *c*-ring are localized to *entry*HC, *a*ARG, and *exit*HC. Outside of these points, *c*-ring likely retains its protonated state to minimize charge exposure and facilitate efficient, unidirectional rotation.

One limitation of this study is the use of coarse-grained model, which limits the resolution for analyzing precise side-chain movements. Since the Cα atom is not located at the tip of the proton-carrying side chain, angular estimates may carry a few degrees of systematic error. However, because this error is uniform across the system, relative comparisons remain valid. Another limitation is that the computational model includes only *a*-subunit and *c*_10_-ring. F_1_ motor, which connects to *c*-ring via the central rotor, and the peripheral stalk are not included. In entire F_o_F_1_, mechanical feedback such as rotational resistance from F_1_ should be present, potentially leading to more asymmetric or complex behavior. Nevertheless, our simplified model has previously been validated against multiple experimental results and all-atom MD simulations, supporting the validity of the observed mechanisms. Thus, the findings obtained here—especially the impact of angular parameters on proton transfer—offer a reliable and meaningful insight into the design principles underlying the rotational efficiency of F_o_ motor.

## Materials & Methods

### Computational System

To simulate the proton-coupled rotational mechanism of *c*_10_-ring, we performed hybrid Monte Carlo/Molecular Dynamics (MC/MD) simulations. The structural model was based on the yeast mitochondrial F_o_ subcomplex from the cryo-EM structure of ATP synthase (PDB ID: 6CP6). The total energy function of the system was defined as:

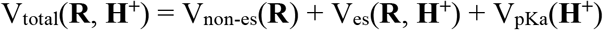

Here, **R** represents all atomic coordinates of the protein, and **H**^**+**^ denotes the protonation states of 12 protonatable residues. V_non-es_(**R**) accounts for all non-electrostatic protein interactions. This term was modeled using the AICG2+ coarse-grained force field, where each residue is represented by a single particle located at the Cα atom. Lipid bilayers and water were treated implicitly. Membrane–protein interactions were modeled through a potential that mimics the hydrophobic environment. V_es_(**R, H**^**+**^) represents electrostatic interactions dependent on both structure and protonation states. This includes standard Coulombic interactions between charged residues and an additional membrane potential term that penalizes deprotonated *c*Glu residues located in the membrane region. The validity of this membrane model has been described in (12). V_pKa_(**H**^**+**^) expresses the energy cost of protonation/deprotonation and depends on environmental factors such as membrane potential (set to 150 mV) and pH (set to 7.0 and 8.0 on the IMS and matrix sides, respectively), as well as the intrinsic pKa values of the residues. In each hybrid simulation step, the protonation states of 12 sites (ten *cE59, a*E223, and *a*E162) were updated using MC sampling. This was followed by a MD phase of 10^5^ MD steps using Langevin dynamics to update atomic positions. A total of 3000 such MC–MD cycles were performed, corresponding to 3.0 × 10^8^ MD steps. For each simulation condition, we conducted ten independent runs with different random seeds. All simulation parameters were consistent with our previous studies.

## Data Availability

The cryo-EM structures used in this study were downloaded from the Protein Data Bank under PDB ID 6CP6. All MD simulations in this study were performed using the CafeMol software. The data were downloaded from https://www.cafemol.org.

## Author Contribution

S.K. and H.N. conceived and designed the project; S.K. developed the simulation code, performed the simulations, analyzed the data, and assembled the figures; S.K. and H.N. discussed the results and all authors were involved in the manuscript writing process.

## Acknowledgements

We would like to thank Ryohei Kobayashi at the University of Tokyo for their support throughout this collaboration. We also thank Kikuyo Ohta for their secretarial assistance.

