## Supplemental Fig. 1 for "Structural Basis for Efficient Fo Motor Rotation Revealed by MCMD simulation and Structural Analysis"

**Classification:** Biological Sciences
**Keywords:** $F_OF_1$ ATPase, molecular dynamics simulation
**Short Title:** Structural Basis of Fo Motor Efficiency

### Supplemental Information

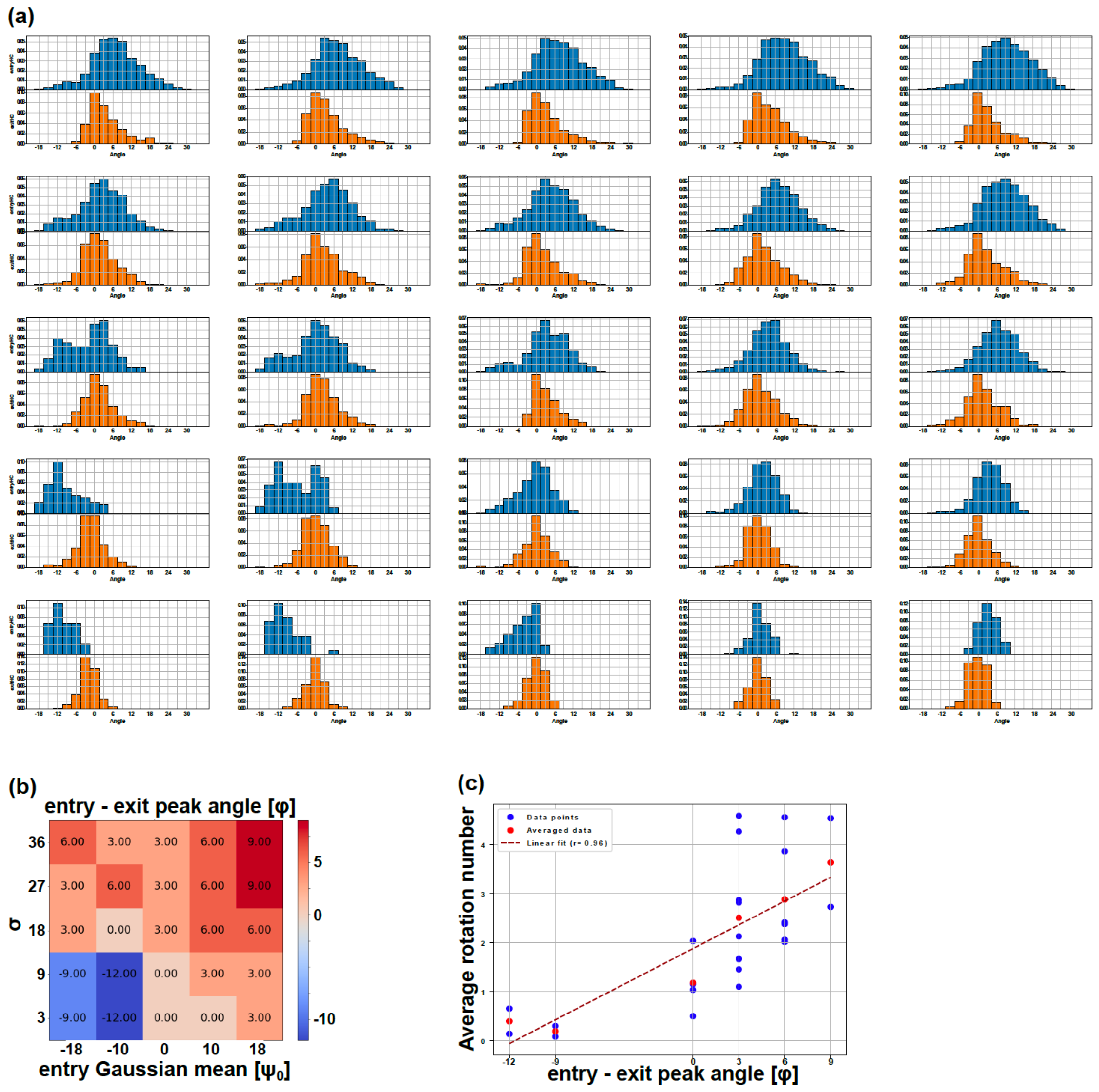


**Figure S1. Angular mapping of proton transfer across diverse σ and $\psi_0$ states. (a)** Histograms of proton transfer angles between *entry*Glu and *c*Glu (top, blue) and between *exit*Glu and *c*Glu (bottom, orange). Layout follows Fig. 3b: rows represent σ= 36, 27, 18, 9, 3; columns represent $\psi_0 = -18, -10, 0, +10, +18$. **(b)** Heatmaps showing the difference in peak angles between Entry and Exit for the combinations of the five $\psi_0$ values and five σ values shown in Fig. 3b. In the heatmaps, values increase from blue to red. **(c)** Average rotation rates plotted against entry–exit peak angle shown in **(b)**. Blue circles indicate values for each (σ, ψ) condition; red circles show averages for each entry-exit peak angle.
